# Optimal sample size calculation for null hypothesis significance tests

**DOI:** 10.1101/540468

**Authors:** Joseph F. Mudge, Jeffrey E. Houlahan

## Abstract

Traditional study design tools for estimating appropriate sample sizes are not consistently used in ecology and can lead to low statistical power to detect biologically relevant effects. We have developed a new approach to estimating optimal sample sizes, requiring only three parameters; a maximum acceptable average of α and β, a critical effect size of minimum biological relevance, and an estimate of the relative costs of Type I vs. Type II errors.This approach can be used to show the general circumstances under which different combinations of critical effect sizes and maximum acceptable combinations of α and β are attainable for different statistical tests. The optimal α sample size estimation approach can require fewer samples than traditional sample size estimation methods when costs of Type I and II errors are assumed to be equal but recommends comparatively more samples for increasingly unequal Type I vs. Type II errors costs. When sampling costs and absolute costs of Type I and II errors are known, optimal sample size estimation can be used to determine the smallest sample size at which the cost of an additional sample outweighs its associated reduction in errors. Optimal sample size estimation constitutes a more flexible and intuitive tool than traditional sample size estimation approaches, given the constraints and unknowns commonly faced by ecologists during study.

## Introduction

Good study design ensures that scientists can make meaningful and appropriate inferences from data. The influence of sample size on inference strength has been well documented in ecology for species richness estimates (Heck et al. 1975), species distributions (Hernandez et al. 2006) and genetic diversity (Leberg 2002). In addition to affecting the precision of parameter estimates, sample size has a major impact on the statistical power of a null hypothesis significance test to detect an effect (Steidl et al. 1997, Jennions and Moller 2003).

Techniques exist for estimating appropriate numbers of samples for most null hypothesis significance tests (Zar 2007). These techniques estimate the minimum sample size needed to achieve the desired power (1-β) to detect the critical effect size, given the expected variability and the chosen α level. Thus, they require specification of α, β, critical effect size and expected variability. All too often these tools go unused (or at least unreported) in ecological research, especially in field-based studies. The reasons for this include that (1) often the replication required to meet accepted power levels is either logistically impractical or prohibitively expensive and sample size may often be restricted by funding, time, space, regulatory, and/or personnel constraints, (2) a number of the parameters in sample size estimation formulas, including critical effect sizes, acceptable α and β levels and even expected variability, are difficult for ecologists to estimate *a prior*i, and (3) ecologists don’t give statistical power the consideration is deserves.

Scientists have reached a somewhat arbitrary consensus on the acceptable α level (i.e.Type I error rate) under the null hypothesis (α = 0.05). Despite this consensus, when expected variability is high and/or biologically relevant effect sizes are small the use of α = 0.05 can require unattainably large numbers of samples. By contrast, there is no consensus on the acceptable β level (i.e.Type II error rate) under the alternate hypothesis. Suggestions for minimum acceptable levels of statistical power include 80%, 90% and 95% (i. e.Type I error rate = Type II error rate). The result is that researchers may forego or ignore sample size estimation and tolerate unreasonably high Type II error probabilities.

Identifying biologically relevant (i.e. critical) effect sizes is difficult and often subjective (Munkittrick et al. 2009). This has led to the standard practice of only evaluating biological significance after having checked for statistical significance (and sometimes erroneously equating statistical significance, or lack thereof, with biological significance) (Martínez-Abraín 2008).

Further, estimation of the expected amount of variability among replicates is also frequently difficult for ecologists. Pilot studies intended to estimate variability levels for sample size estimation can have the same logistical and financial constraints as noted above for the primary study. Further, variability estimates from previously published studies may be unreliable because variability in natural systems often differs among species, locations and times. This has led researchers to invest in methodological techniques that limit variability and thus increase the power to detect effects (Munkittrick et al. 2009).

Mudge et al. (2012) describes an approach for setting optimal α levels for null hypothesis significance tests. This approach minimizes either the combined probabilities or costs of Type I and II errors given a specific combination of critical effect size, relative costs of Type I vs. Type II error and relative prior probabilities of null and alternate hypotheses. Instead of specifying the sample size (among other things) to calculate the smallest possible combination of α and β, the approach can be modified to allow an optimal sample size calculation, given a maximum acceptable combination of α and β.

If the sample size recommendation based on preliminary acceptable error rates and target effect sizes are impractical, researchers would be able to modify either the amount of error they are willing to accept, modify the critical effect size relative to the variability that they wish to be able to detect, or modify the endpoint and/or study design. This approach forces researchers to be explicit about what effects sizes they are targeting and what error rates they are willing to accept (rather than using default Type I error rates and arbitrary Type II error rates). Further, once those decisions are made it allows researchers to identify the minimum number of required samples. This would allow researchers, on the one hand, to avoid doing more sampling than necessary and, on the other, to avoid doing research that doesn’t have the power to detect relevant effect sizes with an acceptable error rate.

Our objectives are to (1) describe how the optimal α approach can be used to calculate optimal (i.e. minimum) sample sizes during study design, (2) show how optimal sample size is affected by the maximum acceptable average of α and β, critical effect size, and relative costs of Type I vs. Type II errors, (3) compare and contrast sample size estimates made using optimal α to traditional sample size estimation approaches, and (4) provide example scenarios that demonstrate how estimating optimal sample sizes could be applied in ecology research.

## Materials and Methods

### Using optimal α to select sample size

Mudge et al. (2012) demonstrated that an optimal alpha can be estimated because for any combination of sample size, critical effect size, relative costs of Type I vs. Type II error and relative prior probabilities of null vs. alternate hypotheses, there exists a single combination of α and β that minimizes the average of Type I and Type II errors. Similarly, an optimal sample size can be estimated because for any combination of desired Type I and II error rates, critical effect size, relative costs of Type I vs. Type II error and relative prior probabilities of null vs. alternate hypotheses, there exists a minimum sample size that can achieve the targeted Type I and Type II error rates. (Note, to simplify the explanation we have assumed that the prior probability of Ho and Ha are equal. However, optimal sample size can be calculated when that assumption isn’t met). Optimal significance levels that minimize combined probabilities of Type I and II errors in null hypothesis tests can be calculated through an iterative process of examining the average of α and β over α values ranging from 0 to 1, to determine the α value that results in the smallest average of α and β. Similarly, for a given critical effect size and relative costs of Type I vs. Type II errors, optimal sample size can be calculated through another iterative process that examines the average of the optimal α and the associated β over a range of sample sizes to iteratively determine the smallest sample size at which the average of α and β remains below the maximum acceptable average of α and β.

Optimal sample sizes will differ for each combination of critical effect size, maximum acceptable average of α and β, relative (or absolute) costs of Type I vs. Type II errors and test type.

### Simulations

To examine how optimal sample size is influenced by other aforementioned parameters, we have begun with a simple, two-tailed, two-sample independent t-test and plotted the optimal sample size at different combinations of critical effect size and maximum acceptable average of α and β, assuming equal costs of Type I vs. Type II error. We then show examples of the optimal sample sizes for the same test-type and the same ranges of critical effect size and maximum acceptable averages of α and β, under four different unequal relative costs of errors to show how optimal sample size changes when either Type I errors are considered to be either 0.1x, 0.5x, 2x or 10x the cost of Type II errors. Then, we return to assuming equal relative costs of Type I and Type II errors and contrast the optimal sample sizes recommended at different combinations of critical effect size and maximum acceptable average of α and β for simple linear correlations and a one-way ANOVA with 2, 4, 6 and 8 treatment groups.

To directly compare sample size recommendations generated using the optimal α approach with sample sizes generated using the traditional sample size estimation technique that requires specifying desired α and β levels, we chose a two-tailed, two-sample t-test over a range of critical effect sizes and relative costs of Type I vs. Type II error and compared differences in sample size recommendations between a standard approach targeting α = 0.05 and β = 0.2 (80% statistical power) and the optimal α approach targeting a maximum acceptable average of α and β of 0.125, the value associated with using α = 0.05 and attempting to achieve 80% statistical power.

### Scenarios

We also present two hypothetical scenarios to demonstrate how optimal sample size calculation can be used in applied ecology, with: (1) Absolute cost of errors not known and relative costs of errors estimated, and (2) Absolute costs of errors estimated.

### Relative costs of errors estimated

The first scenario involved testing for a relationship between the environmental concentration of a chemical and species abundance at different sites, after the chemical has been observed to have a detrimental effect in a laboratory setting. A significant relationship would trigger management actions to reduce chemical inputs to the environment. A Type I error would result in unnecessary management action while a Type II error would result in unmitigated impacts on the species of interest. Absolute costs of Type I and II errors are unknown but two scenarios are presented: (1) Falsely concluding a relationship exists is 0.25 as serious as missing a real relationship (cost of Type I error = 0.25*cost of Type II error, as is loosely implied by using α = 0.05 and 80% statistical power) and (2) detecting a false relationship is equally serious as missing a real relationship. Two maximum averages of α and β are presented: (1) 0.01 where errors, in general, are serious and (2) 0.1 where errors are less serious, e.g. if the species is less valuable and management actions are less expensive. Two critical effect sizes at which management action would be warranted are presented, (1) a relationship where the chemical explains half of the variance in abundance counts (i.e. R^2^ ≥ 0.5) and (2) a relationship where the chemical explains at least one quarter of the variance in abundance counts (i.e. R^2^ ≥ 0.25). The resulting combinations of sample size calculation parameters are summarized in **Error! Reference source not found.**.

**Figure 1:**
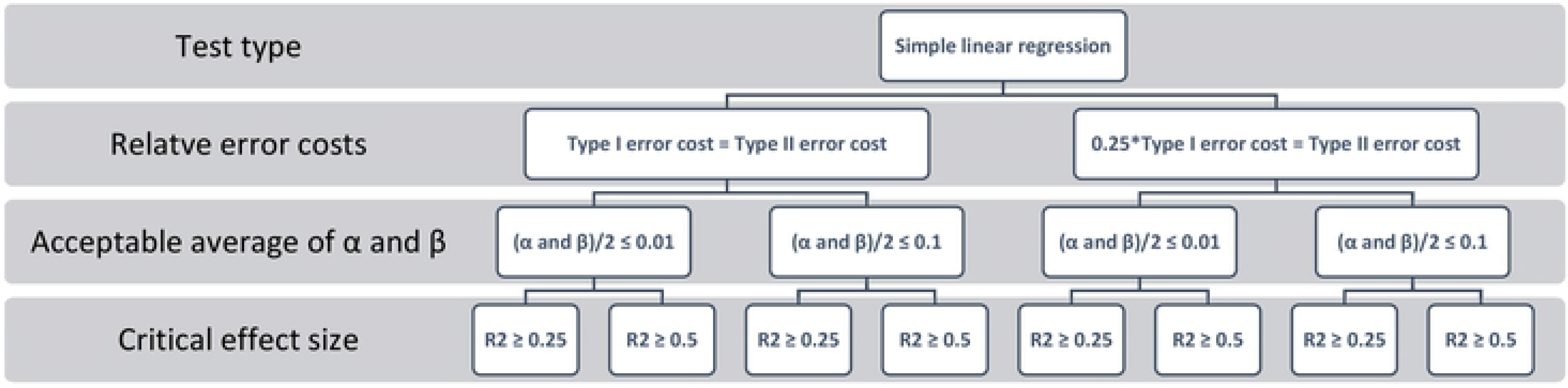
Combinations of relative error costs, acceptable averages of α and β, and critical effect sizes examined for the relative error costs sample size scenario.

### Absolute costs of errors estimated

The second scenario involves monitoring to see if species abundance is significantly greater than the estimate of minimum viable population size. A significant result would signify that the population is well above the minimum viable size and that no management action is required, while a non-significant result would signify that the population is not significantly greater than its minimum viable size and would trigger conservation management actions. In this case, a Type I error would result in managers being unaware that a population is potentially falling below its minimum viable size, while a Type II error would result in unwarranted conservation management actions. Two absolute costs of errors are examined, one where the focal species is economically valuable so a Type I error will cost ten times more than a Type II error (at $10 000 000 vs. $1 000 000 respectively) and another where the cost of conservation is equivalent to the value of the species (at $1 000 000 each). Two potential sample costs were examined, $1000 and $10000 per sample. Also presented are results for two potential critical effect sizes at which proximity to the minimum viable population would trigger conservation management: one and two standard deviations from the minimum viable population size. Note that we don’t set a maximum average error rate because the average error rate is decided by the sample size at which the cost of adding an additional sample would not reduce the costs of an error enough to offset the cost of an additional sample. For example, if the costs of Type I and II errors were each $1,000,000 dollars and the cost of an additional sample was $1,000 we would only add an additional sample if it reduced the average error rate by more than 0.001, The resulting combinations of sample size calculation parameters are summarized in Fig. 2.

**Figure 2:**
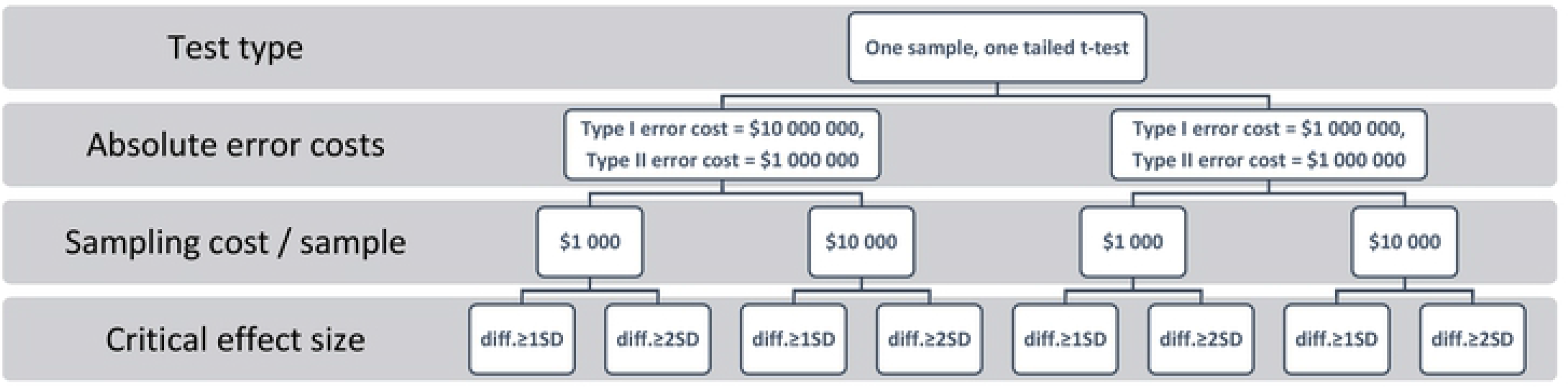
Combinations of absolute error costs, sampling costs, and critical effect sizes examined for the absolute error costs sample size scenario.

## Results

### Simulations

#### t-test: equal costs of errors

Sample size estimates increase as critical effect size and/or the maximum acceptable average of α and β decrease (Fig. 3). For a 2-tailed, 2-sample t-test with equal numbers of samples in each group and equal relative costs of Type I vs. Type II error, fewer than 15 samples are required to detect critical effect sizes ≥ 2 standard deviations of the data at all but the smallest maximum acceptable averages of α and β. By contrast, optimal sample size recommendations are larger than15 for a critical effect size of 1 standard deviation of the data for any maximum acceptable average of α and β > 0.05. More than 30 samples are required to detect effects ≤ 0.5 SD for all maximum acceptable averages of α and β < 0.15. At any given maximum acceptable average of α and β, the relationship between critical effect size and optimal sample size is nonlinear, requiring increasingly large increases in sample size as critical effect size decreases. For example, if we set the maximum acceptable average of α and β at 0.125, halving the critical effect size from 1.0 to 0.5 SD more than triples the optimal sample size (Fig. 4).

**Figure 3:**
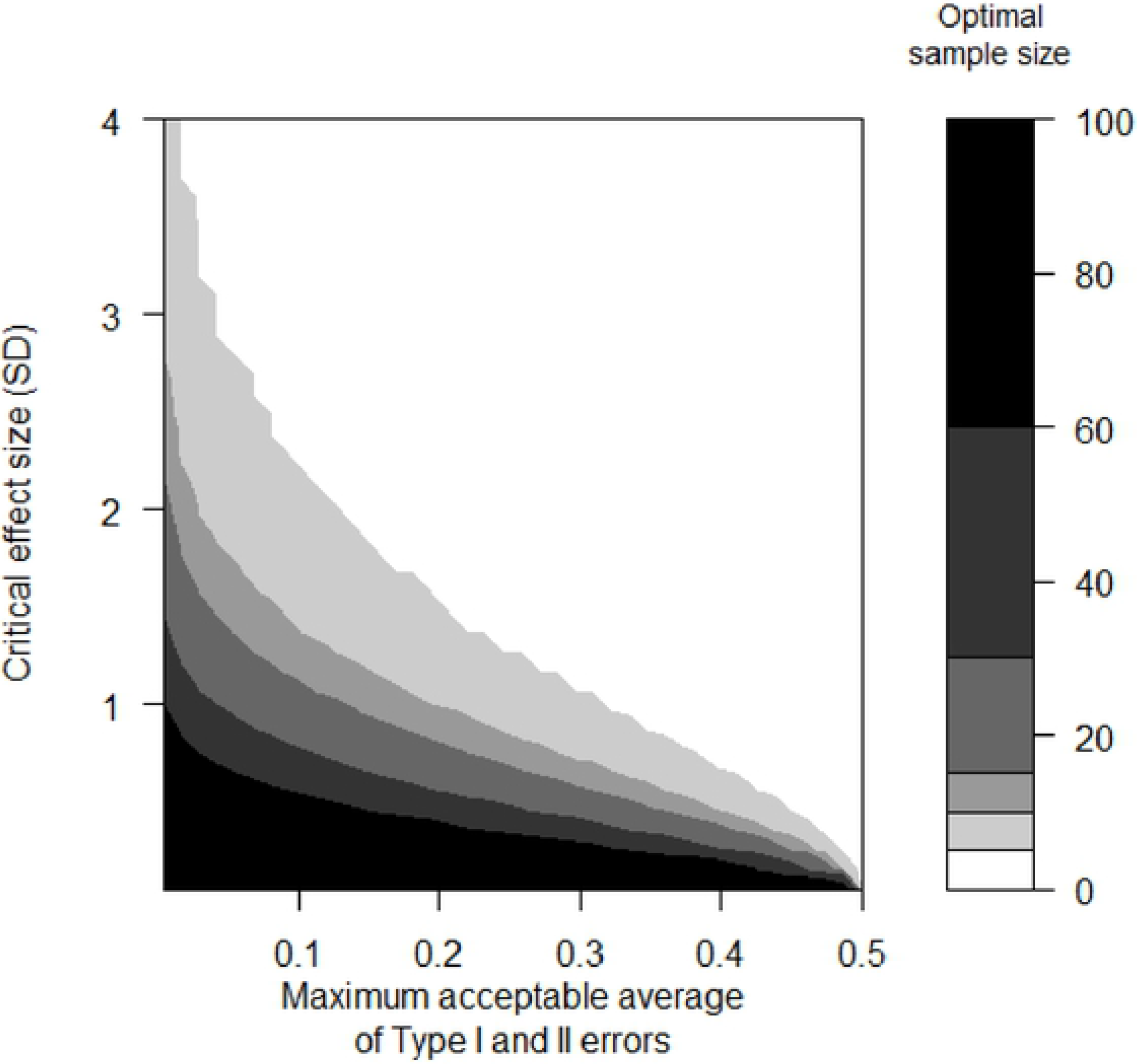
The minimum number of samples in each group that are required to achieve the desired maximum acceptable average of Type I and Type II error rates over a range of critical effect sizes (expressed in standard deviation units), for a 2-tailed, 2-sample t-test with an equal number of samples in each group and equal relative costs of Type I vs. Type II error.

**Figure 4:**
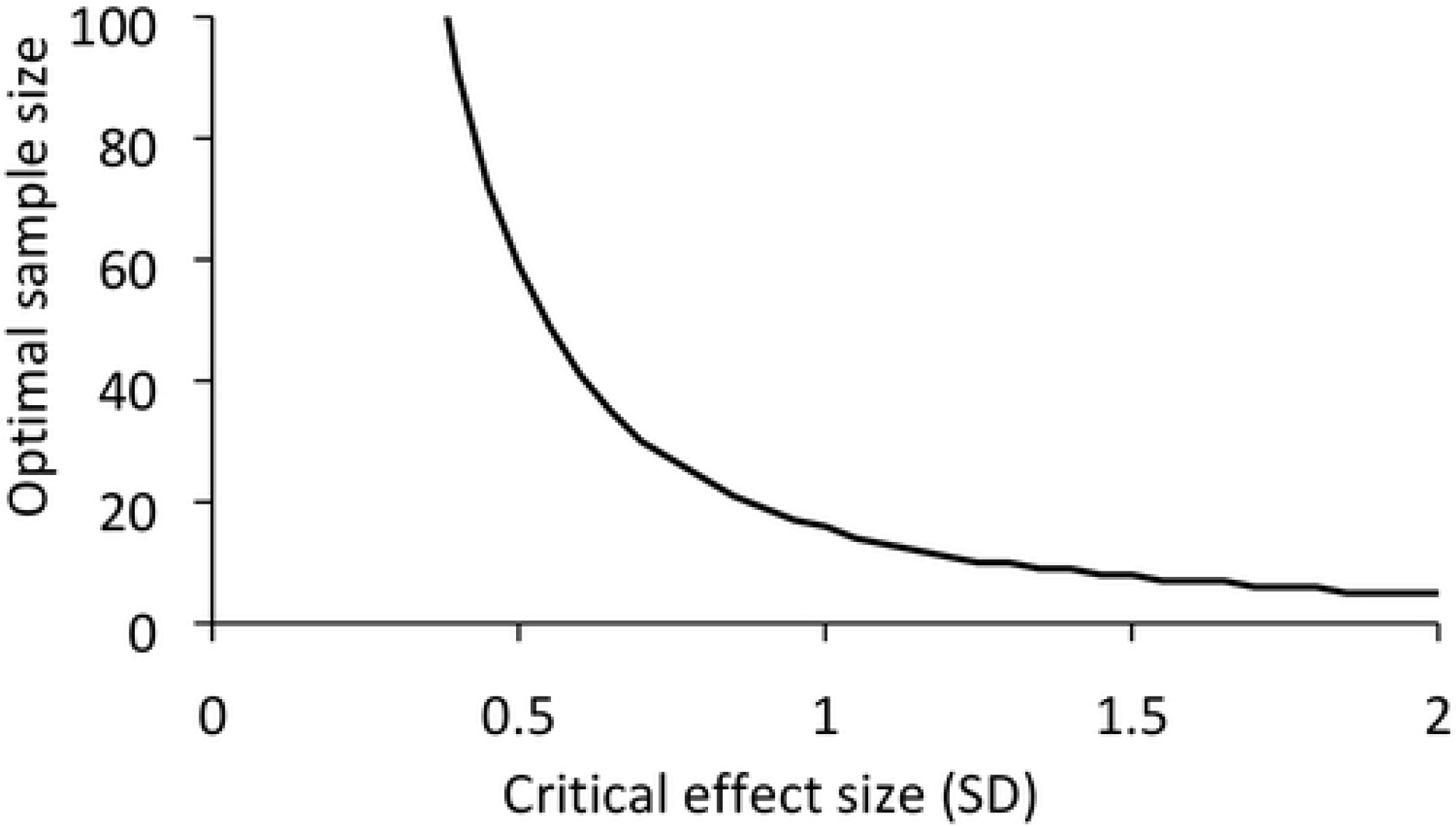
The influence of the choice of critical effect size (expressed in standard deviation units) on the optimal sample size for a two-tailed, two-sample t-test, assuming equal relative costs of Type I and II errors and holding the maximum acceptable average of α and β at 0.125.

#### t-test: unequal costs of errors

When relative costs of Type I and Type II error are unequal, larger sample sizes are required to detect the same combinations of critical effect size and maximum acceptable average of α and β, relative to those recommended for equal relative costs of Type I and II errors (Fig. 5) and the greater the deviation from equality of the relative costs of Type I and II errors, the greater the increase in required sample size.However, the influence of relative error costs on sample size recommendation is much smaller than the influence of either critical effect size or maximum acceptable average of α and β. There are also marginal differences in recommended sample size depending on which error type is more costly.

**Figure 5:**
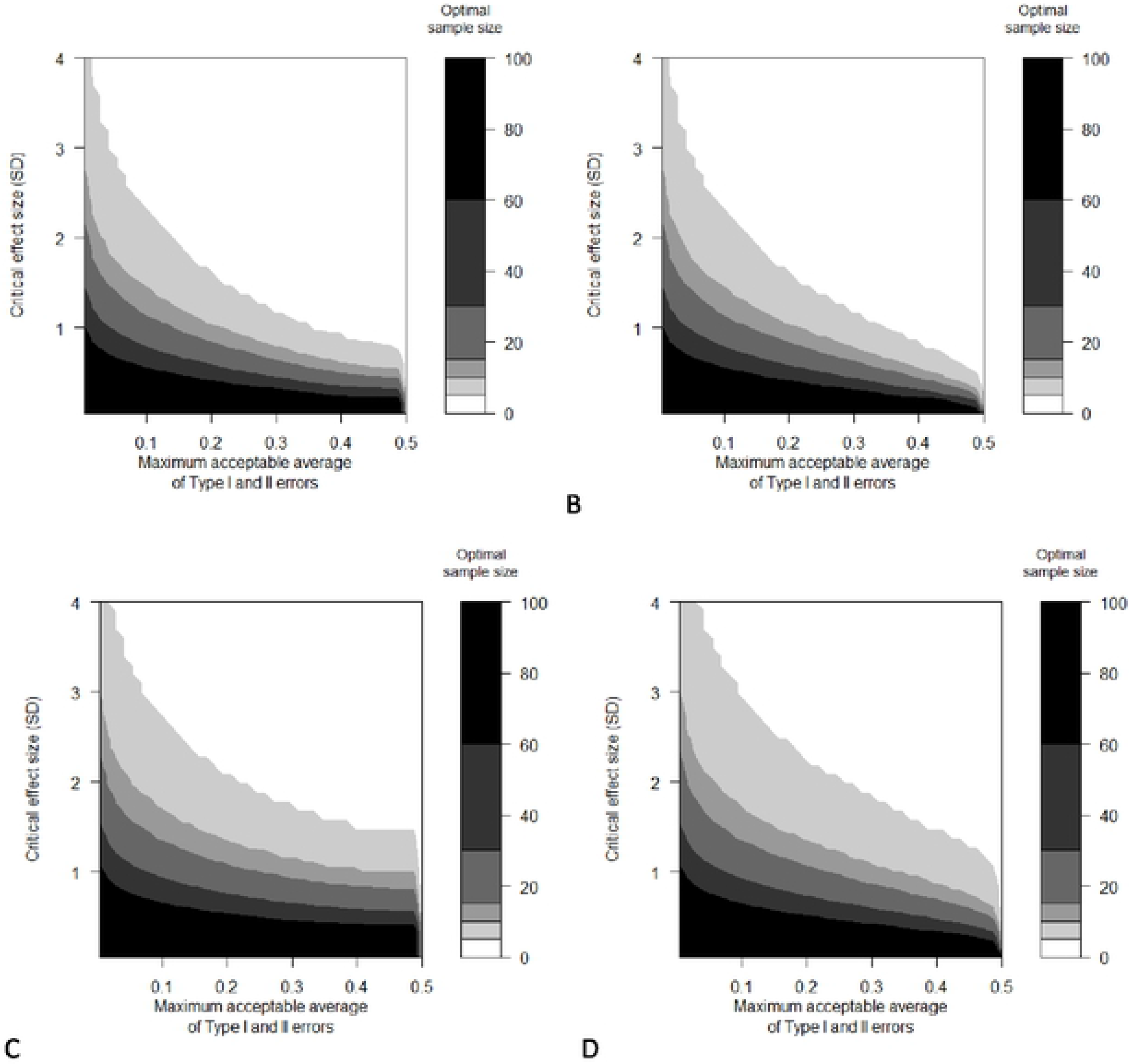
The minimum number of samples in each group that are required to achieve the desired maximum acceptable average of Type I and Type II error rates over a range of critical effect sizes (expressed as standard deviation units), for a 2-tailed, 2-sample t-test with an equal number of samples in each group and (A) Type I errors considered half as serious as Type II errors, (B) Type I errors considered twice as serious as Type II errors (C) Type I errors considered 10 times less serious than Type II errors and (D) Type I errors considered 10 times more serious than Type II errors.

### Correlation and ANOVA tests

The influences of critical effect size and maximum acceptable average of α and β on sample size recommendation differ among test types (Fig. 6, Fig. 7). For simple linear correlation tests (Fig. 6), sample sizes smaller than 10 are rarely informative, as these small sample sizes are only recommended when either the critical effect size is ≥ r = 0.7 (R^2^ = 0.5) and/or the maximum acceptable average of α and β is ≥ 0.2. Sample sizes approaching 30 can be more informative, with these sample sizes being recommended when either the critical effect size is r = 0.5 (R^2^ = 0.25) and/or the maximum acceptable average of α and β is ≥ 0.125. In ANOVA study designs (Fig. 7), increasing the number of levels of a factor decreases the number of replicates required within each level of the factor. However, this decrease in the number of replicates required within each factor level comes at the expense of an even greater increase in the total number of replicates required among all factor levels, such that using more replicates within fewer factor levels is more efficient than using fewer replicates within more factor levels. The total number of samples in an ANOVA design required to achieve a maximum acceptable average of α and β of 0.05 for a critical effect size of f^2^ = 1 is 16 with 2 groups (8 replicates * 2 groups), 24 with 4 groups (6 replicates * 4 groups), 30 with 6 groups (5 replicates * 6 groups), and 32 with 8 groups (4 replicates * 8 groups).

**Figure 6:**
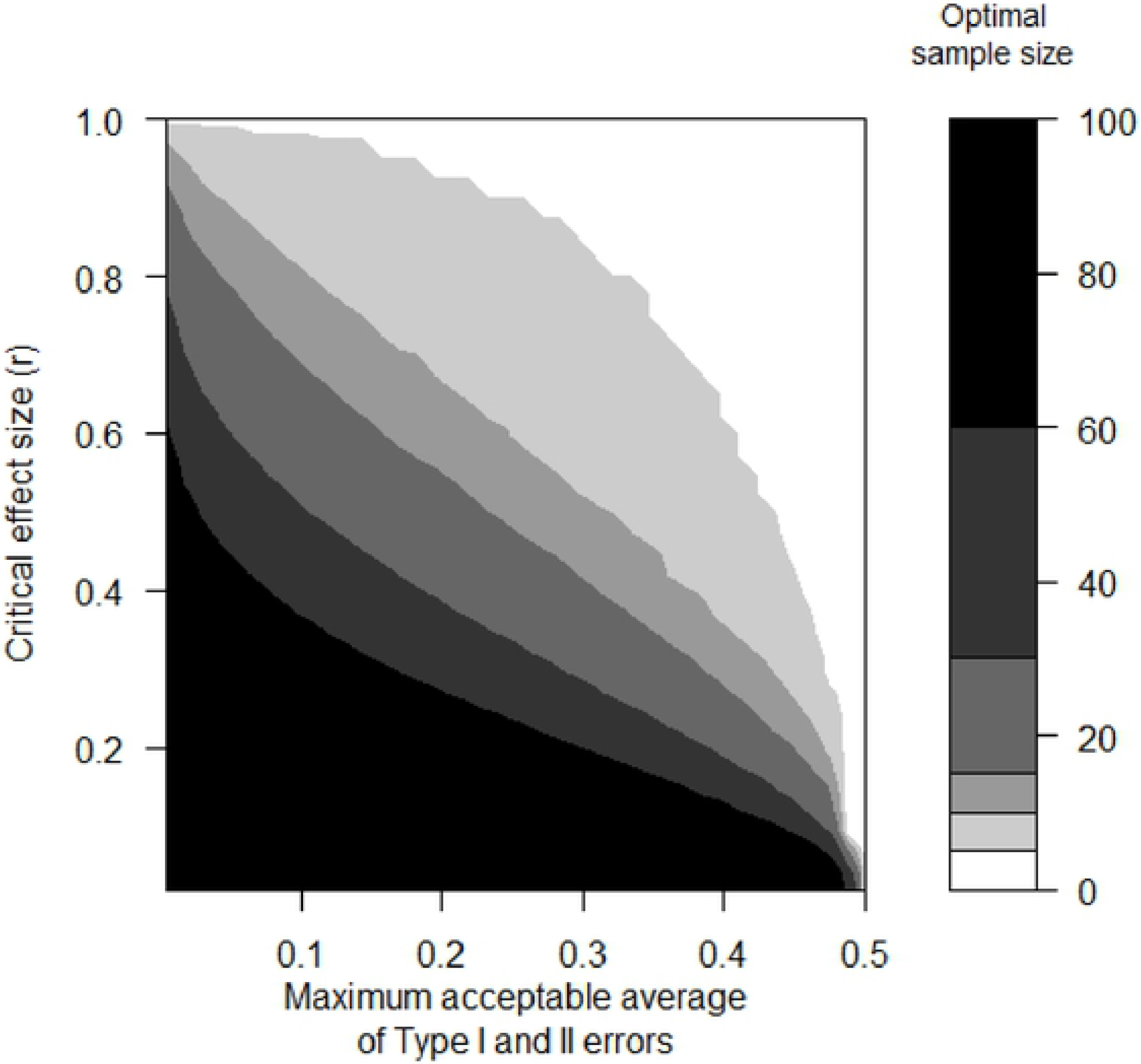
The minimum number of samples that are required to achieve the desired maximum acceptable average of Type I and Type II error rates over a range of critical effect sizes (expressed as a correlation coefficient), for a 2-tailed simple linear correlation test with equal relative costs of Type I vs. Type II error.

**Figure 7:**
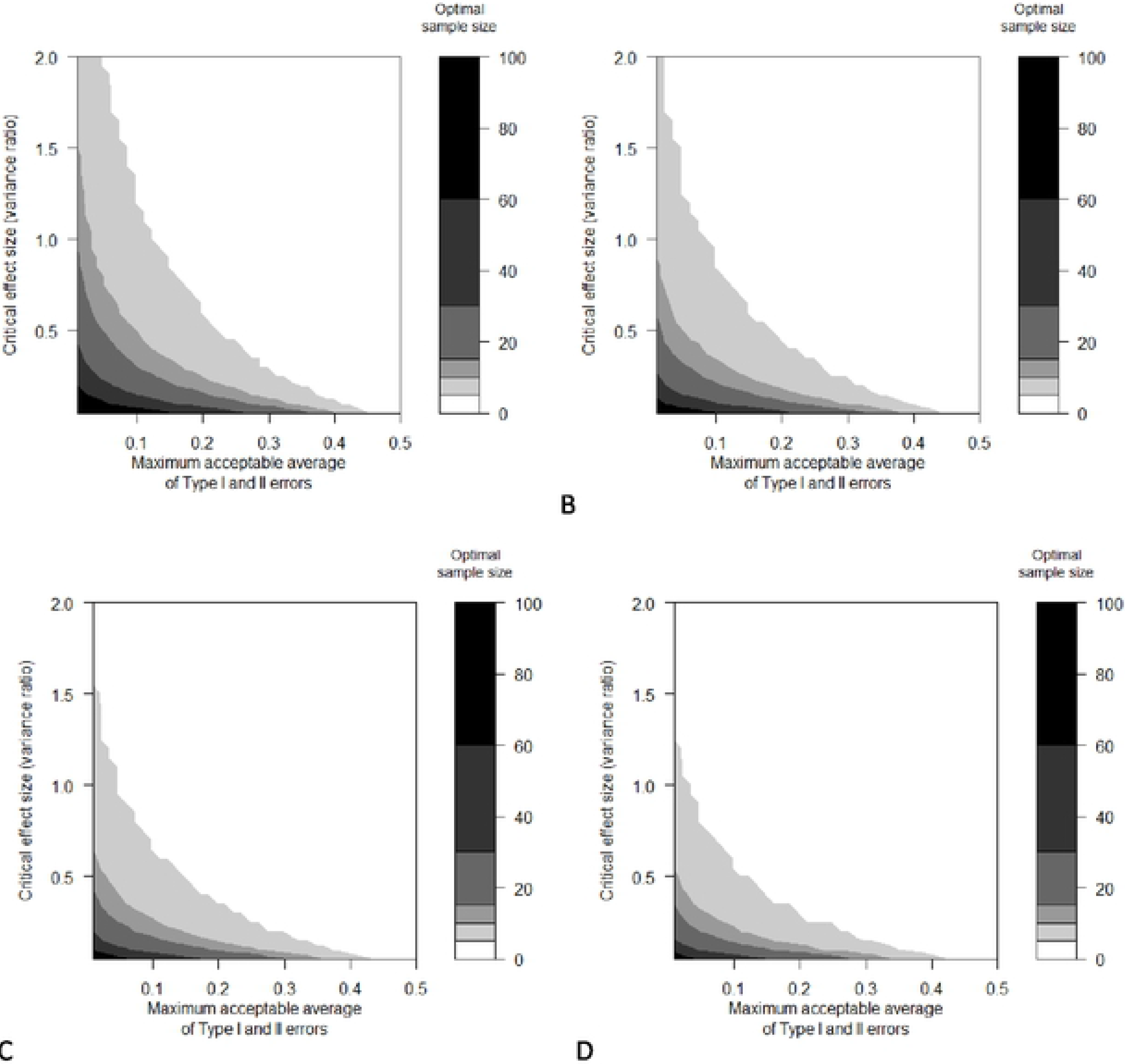
The minimum number of replicates in each group that are required to achieve the desired maximum acceptable average of Type I and Type II error rates over a range of critical effect sizes (expressed as a ratio of among-group variance to within-group variance), for a one-way ANOVA with an equal number of replicates in each group, equal relative costs of Type I vs. Type II error and (A) 2 treatment groups, (B) 4 treatment groups (C) 6 treatment groups and (D) 8 treatment groups.

### t-test: comparison between optimal α and traditional sample size recommendations

Optimal sample sizes increase as the critical effect size decreases and as relative costs of Type I and II errors deviate from equality. The traditional approach of designing t-test studies to achieve 80% statistical power with an α = 0.05 also results in larger sample size recommendations as critical effect size decreases, however this approach does not recommend adjusting sample sizes when relative costs of Type I and II errors differ. When relative costs of Type I and II errors are equal, the optimal α sample size estimation approach can achieve the same maximum acceptable average of α and β at the same effect size using fewer samples than the traditional sample size estimation approach using α = 0.05 and 80% statistical power (Fig. 8). As relative costs of Type I vs. Type II errors become more unequal, the sample sizes recommended by the optimal α approach increase to the point where they surpass those recommended by the traditional sample size estimation approach using α = 0.05 and 80% statistical power. Optimal α sample size recommendations are larger than those recommended by traditional sample size estimation approach for highly unequal relative costs of Type I vs. Type II errors, however when relative costs of Type I and II errors are highly unequal, the practice of using α = 0.05 and 80% statistical power becomes less and less justifiable.

**Figure 8:**
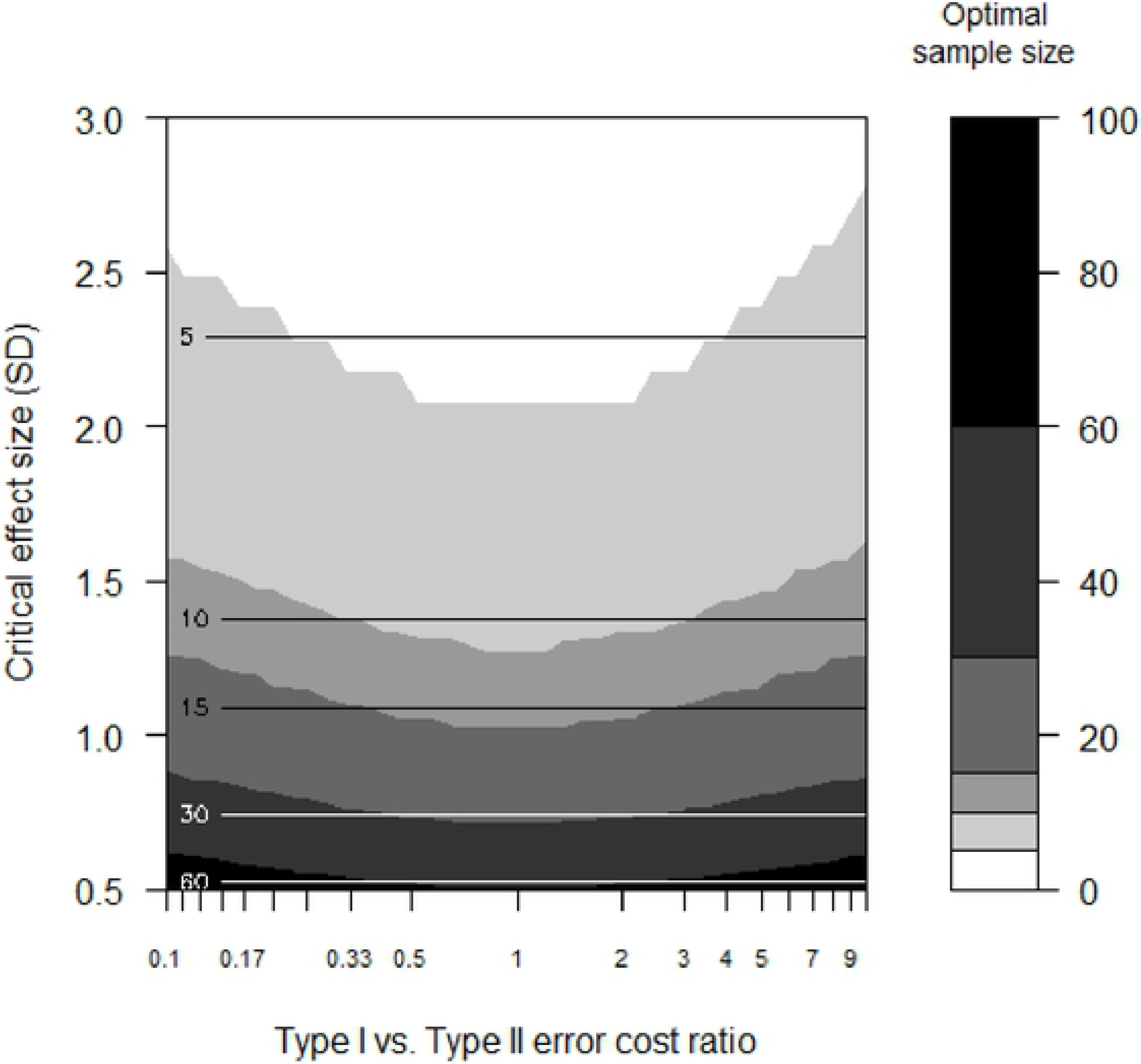
Comparison of the sample size recommendations between the optimal α approach (shaded areas) and the traditional sample size estimation approach (horizontal lines), over a range of critical effect sizes (expressed in standard deviation units) and relative costs of Type I vs. Type II errors, for a two-tailed, two-sample t-test using a maximum acceptable average of α and β of 0.125 for the optimal α approach and α =0.05 and β =0.2 for the traditional sample size estimation approach.

### Scenarios

#### Absolute costs of errors unknown

Optimal sample size recommendations ranged from 14 to 89, depending on the combination of Type I vs. Type II error cost ratio, maximum acceptable average of α and β, and critical effect size (Table 1). Decreasing the Type I vs. Type II error cost from 1 to 0.25 resulted in small increases in sample size recommendations of 2 to 6 samples. A 10-fold decrease in the maximum acceptable average of α and β, and a 2-fold decrease in critical effect size each resulted in a greater than 2-fold increase in the sample size recommendation.

**Table 1:** Optimal sample sizes that achieve a maximum acceptable average of α and β for a simple linear regression under different relative costs of Type I vs. Type II error, maximum acceptable averages of α and β and critical effect sizes, with associated α and β levels for the optimal sample size.

| Type I vs. Type II error cost ratio | Maximum acceptable average of $\alpha$ and $\beta$ | Critical effect size ($R^2$) | Optimal sample size | Optimal $\alpha$ | Optimal $\beta$ |
| --- | --- | --- | --- | --- | --- |
| 1 | 0.01 | 0.5 | 34 | 0.010 | 0.010 |
| 1 | 0.01 | 0.25 | 83 | 0.009 | 0.010 |
| 1 | 0.1 | 0.5 | 14 | 0.091 | 0.103 |
| 1 | 0.1 | 0.25 | 31 | 0.091 | 0.108 |
| 0.25 | 0.01 | 0.5 | 37 | 0.015 | 0.003 |
| 0.25 | 0.01 | 0.25 | 89 | 0.016 | 0.004 |
| 0.25 | 0.1 | 0.5 | 16 | 0.166 | 0.033 |
| 0.25 | 0.1 | 0.25 | 37 | 0.164 | 0.033 |

#### Absolute costs estimated

Optimal sample size recommendations ranged from 6 to 36, depending on the combination of Type I error cost, cost per sample, and critical effect size (Table 2). Increasing the cost of Type I errors from $1 000 000 to $10 000 000 resulted in sample size recommendations increasing by 4-10 samples. A 10-fold decrease in the cost per sample and a 2-fold decrease in the critical effect size each had large impacts on recommended sample size, each increasing the sample size recommendation more than 2-fold. Total sampling costs were lowest when costs of errors were equal, when the critical effect size was large and, as expected, when costs per sample were low.

## Discussion

### Comparison between optimal and traditional sample size estimation

Ecologists must consider the effect size that would be important to detect, the relative seriousness of different potential errors and how much error they would be willing to accept as central to good study design. The optimal α approach can directly incorporate each of these important considerations in the calculation of either the minimum sample size that achieves the maximum acceptable average of α and β, or the minimum sample size at which the cost of an additional sample outweighs the reduction in errors associated with the additional sample. By contrast, the traditional sample size estimation approach of calculating the minimum sample size required to achieve a specific β level fails to minimize the number of samples required.

**Table 2:** Optimal sample sizes at which the cost of an additional sample exceeds the reduction in error costs for a one-tailed, one-sample t-test under different absolute error costs, sample costs and critical effect sizes, with associated α and β levels and the total sampling cost for the optimal sample size.

| Cost of Type I error (\$) | Cost of Type II error (\$) | Cost per sample (\$) | Critical effect size (Cohen's d) | Optimal sample size | Optimal $\alpha$ | Optimal $\beta$ | Total sampling cost (\$) |
| --- | --- | --- | --- | --- | --- | --- | --- |
| 1 000 000 | 1 000 000 | 1 000 | 2 | 12 | 0.0033 | 0.0017 | 12 000 |
| 1 000 000 | 1 000 000 | 1 000 | 1 | 26 | 0.0092 | 0.0072 | 26 000 |
| 1 000 000 | 1 000 000 | 10 000 | 2 | 6 | 0.0341 | 0.0157 | 60 000 |
| 1 000 000 | 1 000 000 | 10 000 | 1 | 10 | 0.0791 | 0.0587 | 100 000 |
| 10 000 000 | 1 000 000 | 1 000 | 2 | 16 | 0.0003 | 0.0019 | 16 000 |
| 10 000 000 | 1 000 000 | 1 000 | 1 | 36 | 0.0008 | 0.0080 | 36 000 |
| 10 000 000 | 1 000 000 | 10 000 | 2 | 10 | 0.0027 | 0.0180 | 100 000 |
| 10 000 000 | 1 000 000 | 10 000 | 1 | 18 | 0.0070 | 0.0809 | 180 000 |

### Critical effect sizes and sample size estimation

The critical effect size chosen has a strong influence on the sample size recommendation. In ecology, the small sample sizes that are frequently found in both lab and field experiments (i.e. less than 10 samples) are only optimal for t-tests when critical effect sizes are moderate to large (≥1.5 SD) (if we assume ((α+β)/2 = 0.125, which occurs with α = 0.05 and 80% statistical power). Many researchers may be unaware that they are making the implicit assumption that small to moderate effects are not important. The larger sample sizes that are more frequently found in observational studies (10-30 samples) have the capability to detect smaller effects in terms relative to the within-group standard deviation (≥1 SD), however the within-group standard deviation is often larger for observational studies than in comparatively more controlled laboratory (or even field) experiments, meaning that a 1 SD effect represents a larger absolute effect for observational studies than for tightly controlled experiments. Critical effect sizes of 0.5 SD or smaller require sample sizes that are rarely achieved in ecological research (> 60 samples). In these situations, ecologists need to either 1) accept only being able to detect moderate to large effects, 2) tolerate larger averages of α and β or 3) use more samples than is currently typical for ecological research.

### Relative costs of Type I vs. II errors and sample size calculation

The optimal α sample size approach shows that a few more samples should be used when one type of error is more serious than the other, relative to the number of samples recommended when Type I and II errors are equally serious. Relative costs of error are rarely addressed in ecological research and even less frequently are they incorporated into study design. The traditional approach to sample size estimation accepts whatever cost-ratio results in α = 0.05 and β = 0.2 at the researcher’s chosen critical effect size. Note, that this does not correspond to a Type I vs. Type II error cost ratio of 4, as assumed by Mapstone (1995), since the error probability ratio and the error cost ratio are not the same (Mudge et al. 2012). Mapstone (1995) recommended setting α and β so as to make β/α ratio equal to the Type I / Type II error cost ratio both for sample size estimation and statistical analysis, however the flaw in this approach can be most easily illustrated when costs of Type I and II errors are equal. It may not be intuitively obvious but, when α and β are set to be equal to correspond with equal costs of errors, the outcome of shifting α and β away from equality will virtually always yield a smaller average of α and β, and trading an increase in one error type for a greater corresponding reduction in the other error type is always preferable when Type I and II errors have equal costs.

Another approach to incorporating relative costs of errors into study design has been proposed by Field et al. (2004). Instead of attempting to minimize the probabilities or costs of errors, this approach can be used to estimate sample sizes that minimize the total cost of monitoring and management according to the probabilities and costs of errors. This approach to sample size estimation with a single goal of minimizing monitoring and management costs can result in recommending a sample size of zero and essentially just assuming there either is an effect or no effect when sampling costs are high and there is a large discrepancy between the costs of Type I and II errors. While this approach may be valuable for applied environmental management, it has received little attention in ecological research. In purely scientific research the ultimate goal is accuracy of results, not cost-effectiveness so that, having low combined probabilities of Type I and II errors is the main priority, and balancing Type I and II errors appropriately according to their respective costs is of secondary concern.

### Differences among statistical test-types

The relationship between the maximum acceptable average of α and β, critical effect size, and optimal sample size does not remain consistent among statistical techniques. For correlation/regression tests, the sample size recommendation curves on the graph of critical effect size vs. maximum acceptable average of α and β switch from curving outward away from the origin to curving inward toward the origin at between 10 and 15 samples (Fig. 6). This means that below 10 to 15 samples, correlation/regression tests are particularly ineffective for testing null hypotheses as they are forced to choose between a very large critical effect size (for which null hypothesis testing becomes increasingly unnecessary), or a very high maximum acceptable average of α and β (or both). For ANOVA, the sample size recommendation curves on the graph of critical effect size vs. maximum acceptable average of α and β have the same general shape as for a t-test, with the key difference being that the curves shift downward as the number of ANOVA factor levels increases, indicating that it takes fewer replicates per factor level to detect effects of a given effect size as the number of factor levels increases. However, the total number of samples required to detect a particular effect size will always be less with fewer factor levels. This means that to minimize sample sizes in ANOVA designs, one should use fewer factor levels whenever possible. Other arguments for fewer factor levels in ANOVA designs are that ANOVA results can be less informative as the number of factor levels increases (i.e. a result of significant variability among many group means may not be surprising or useful for a large number of groups), and the reliability of post-hoc pairwise comparison tests decreases as the number of replicates within factor levels decreases.

### Maximum acceptable average of α and β and sample size estimation

What makes an appropriate maximum acceptable average of α and β? This is a question that ought to concern researchers, funding/granting agencies, regulatory agencies and academic publishers, but that is rarely dealt with directly. The most common combination of α and β used under the traditional sample size estimation approach is α = 0.05 and β = 0.2 (i.e. 80% statistical power), which yields an implied maximum acceptable average of α and β of 0.125. All else being equal, a low average of α and β is always better, as the average α and β is an excellent indicator of the quality of experimental design. Higher maximum acceptable averages of α and β than 0.125 may sometimes be appropriate, especially when samples are very expensive or difficult to collect. One example of this is whole-system studies, which provide a level of realism that is rarely attainable in smaller-scale studies with larger sample sizes. It is important to note that the maximum upper threshold for maximum acceptable averages of α and β would be anything approaching 0.5, the probability of reaching the correct conclusion using a coin toss. When samples are cheap and easy to collect and/or when errors are very serious, a smaller average of α and β than 0.125 may be warranted. Although the maximum acceptable average of α and β is something that is best evaluated on a study-specific basis, threshold maximum acceptable averages of α and β may sometimes be useful as one criterion for evaluation of grant application/renewal or as a component of the evaluation of manuscripts submitted for publication in peer-reviewed scholarly journals. A suitable default maximum acceptable average of α and β could be 0.05, simply because we have become accustomed to the 5% value for Type I errors and confidence intervals. This would result in the default being ‘tradition’ but tradition used in a sensible way. Researchers should, however, start thinking more rigorously about what average of α and β is acceptable, especially because using 0.05 as the default maximum acceptable average of α and β is a more rigorous experimental design than the traditional approach of using α = 0.05 and β = 0.2, and may require more samples to achieve.

### Conclusion

The optimal α approach can be used to calculate optimal sample sizes during study design, recommending few samples when critical effect sizes are large, maximum acceptable averages of α and β are large and relative costs of Type I vs. Type II errors are equal, and many samples at small critical effect sizes and maximum acceptable averages of α and β and unequal relative costs of Type I vs. Type II errors. Holding the maximum acceptable average of α and β constant, the optimal α approach recommended fewer samples than the traditional sample size estimation approach when relative costs of Type I and II errors were approximately equal, and more samples when relative costs were highly unequal. This is because the traditional approach is based on a fundamental assumption that relative costs aren’t equal (i.e. α = 0.05 and β = 0.20) so when the relative costs of error are equal, it is not as efficient as optimal α. When the true relative costs are unequal the traditional approach recommends fewer samples but doesn’t incorporate the true relative costs of error and so will not select the optimal number of samples. The flexibility of optimal sample size estimation allows for sample size estimates that achieve a desired maximum acceptable average of α and β given relative costs of Type I and II errors. However, when absolute Type I and II error costs are known it also allows for sample size estimates that identify when the cost of an additional sample would exceed the reduction in overall error costs associated with adding the extra sample.

#### A brief guide to optimal sample size calculation

The first step in calculating sample sizes using the optimal α approach is to choose values for the parameters on which the optimal sample size calculation will be based: the maximum acceptable average of α and β, the relative costs of Type I and II errors, and the critical effect size. Each of these parameters deserve rigorous consideration, however there are some sensible default values that may be used in the absence of a strong rationale for any particular value. Using 0.05 for the maximum acceptable average of α and β would be consistent with the 5% standard with which we have become accustomed to using for Type I errors and confidence intervals, while using 0.125 would be consistent with the common desire to achieve 80% statistical power at α 0.05. For the relative costs of Type I and II errors, the most appropriate default value (especially for purely scientific research) is to assume equal costs of Type I and II errors, in favour of minimizing their combined probabilities. Although differences among research questions precludes the potential for a default critical effect size that can be used for any study design, see Nakagawa and Cuthill (2007) and Munkittrick et al. (2009) for discussions of how to choose critical effect sizes that are associated with biological relevance.

Optimal sample sizes can be calculated for t-tests, ANOVA and regression/correlations using code available online from Mudge et al. 2012.

## Authors Contributions

JM and JH conceived the ideas. JM designed the methodology and simulations and took the lead on writing. JM and JH contributed to drafts, interpretation of simulation results and approved the final version.

## Acknowledgements

We are grateful for Natural Sciences and Engineering Research Council funding.

